# Improving Neural Networks for Genotype-Phenotype Prediction Using Published Summary Statistics

**DOI:** 10.1101/2021.11.09.467937

**Authors:** Tianyu Cui, Khaoula El Mekkaoui, Aki Havulinna, Pekka Marttinen, Samuel Kaski

## Abstract

Phenotype prediction is a necessity in numerous applications in genetics. However, when the size of the individual-level data of the cohort of interest is small, statistical learning algorithms, from linear regression to neural networks, usually fail due to insufficient data. Fortunately, summary statistics from genome-wide association studies (GWAS) on other large cohorts are often publicly available. In this work, we propose a new regularization method, namely, *main effect prior (MEP)*, for making use of GWAS summary statistics from external datasets. The main effect prior is generally applicable for machine learning algorithms, such as neural networks and linear regression. With simulation and real-world experiments, we show empirically that MEP improves the prediction performance on both homogeneous and heterogeneous datasets. Moreover, deep neural networks with MEP outperform standard baselines even when the training set is small.

## 1 Introduction

Genotype-based phenotype prediction is central in various genetics applications. For example, accurate prediction of human complex traits and diseases from genotype has shown great promise in precision medicine and healthcare [1,7,47]. In genomic selection, predicting the grain yield with genotype and environment factors is revolutionizing plant breeding and increasing food production [34,38,39]. In bacterial genomics, predicting phenotypes such as antibiotic resistance has become a fundamental task to understand bacteria [27,40].

Existing approaches for phenotype prediction can be divided into two categories, depending on whether the modeler has access to individual-level data. Classical statistical genetics approaches, such as BLUP [3,17,33] and its extensions [16,21,44], demonstrate impressive predictive performance when individual-level data are available. Moreover, standard machine learning methods are becoming appealing alternatives because fewer assumptions are made about the genetic mechanism underlying the traits of interest, ranging from linear models, e.g., Lasso [12,28], to complex models, e.g., neural networks [2,18,38,39,41,55], that capture nonlinear main effects and feature interactions. However, due to the few assumptions made by deep neural networks, suitable inductive biases [37,52], such as neural network architectures and regularizations, are needed to learn the optimal function, especially for small data. On the other hand, accessing individual-level genomic data is cumbersome due to the enormous volume of the data and often also restricted due to privacy considerations such as the GDPR [50], which recently have made the summary statistics approach attractive: Models are learned using summary statistics from genome-wide association studies (GWAS) without access to individual-level data [15,31,57]. These methods build on summary statistics, including the estimated main effect and standard error of each single-nucleotide polymorphism (SNP) as well as the estimated linkage disequilibrium (LD) matrix, which are generally easier to share than individual-level datasets.

The accuracy of predictive models trained with individual-level data or summary statistics data for several traits has already been promising [31,32] with large biobank datasets of target cohorts. However, a more realistic situation is that users only have a *small* number of individual-level data for the cohort of interest, in which case neural networks and even linear models may fail due to insufficient data. Fortunately, the GWAS summary statistics of the same traits may be publicly available for a different cohort with a large amount of data. Therefore, the question of how to leverage the external publicly available summary statistics to improve phenotype prediction with complex models on small datasets is an engaging research question with various applications.

In this paper, we focus on predicting continuous human phenotypes, such as metabolites, with genotype data on a small target cohort but with access to the GWAS summary statistics from an external dataset, as shown in Figure 1. We propose a novel regularization term that incorporates prior knowledge about the main effect of each SNP from the external GWAS summary statistics, and the regularization can be widely applied to any optimization-based machine learning method, such as linear models and neural networks. We demonstrate that the *main effect prior (MEP)* consistently improves all models, and neural networks with MEP outperform existing approaches even when the training dataset is small.

**Fig. 1.**
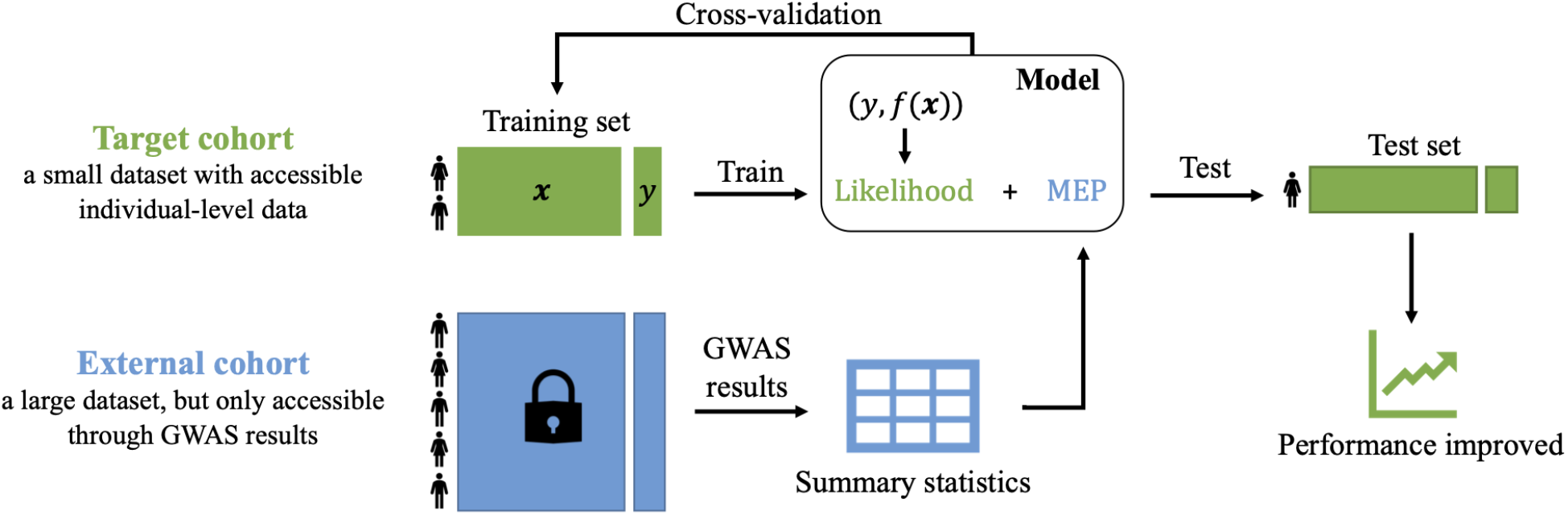
Overview. Prediction accuracy of a model on the target cohort, with few individual-level training data, is improved by incorporating the GWAS summary statistics from an external cohort through our main effect prior (MEP). This does not require accessing the individual-level data of the external cohort.

## 2 Background

### 2.1 Neural networks and regularization

Recent advances in neural networks have demonstrated strong predictive performance with large training datasets in various fields [13,25]. However, neural networks are prone to overfitting when encountered with small and noisy datasets, such as genotype-phenotype datasets. A number of regularization techniques, e.g., weight decay [26] and Lasso neural network [29], have been applied to constrain the flexibility of neural networks; They are all equivalent to zero-mean priors from a Bayesian perspective. More sophisticated regularization techniques have been proposed to incorporate extra domain knowledge into the model beyond sparsity [43]. For example, when the importance score of each feature is known *a priori*, attribution priors could be used to regularize the feature importance in the model, and methods such as DeepSHAP [48] and Contextural Decomposition [42] agree with the corresponding prior. While this way of regularizing the model has improved performance in specific problem domains, interpretation of the results requires *local interpretations* [51], i.e., the feature importance specific to each data point, assigned by humans, instead of *global interpretations*, such as main effects. A further complication is that feature importance scores in the deep learning community have a very different meaning from the main effects that are typically considered in a GWAS. Thus, new ways of regularizing the model are needed for neural networks to incorporate external GWAS results.

### 2.2 Bayesian approaches to incorporate domain knowledge

Priors on the proportion of variance explained (PVE) have been proposed for linear regression [56] and neural networks [11] to improve feature selection and prediction in genetics. Building of informative priors for neural networks has also been studied in the function space. For example, Gaussian processes [14,45] have been proposed as a way of defining functional priors to encode rich functional structures with specific kernels. Bayesian approaches have the capability of marginalizing over all possible weights and capturing the predictive uncertainty [52], with the cost of increased computational complexity, because approximate inference methods are usually needed for neural networks. In this paper, we focus on the regularization-based approaches and leave the Bayesian extensions for future work.

## 3 Methods

### 3.1 Neural networks with residual connections

Let 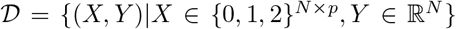 denote a training dataset with *N* individuals, *p* SNPs, and one phenotype variable, where 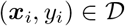 represents the SNPs coded as 0,1 or 2 and phenotype of the *i*th individual. A neural network (NN) is a function mapping inputs to outputs by aggregating predictions of simple but non-linear functions. Selecting a suitable architecture for the NN is critical because it reflects the inductive biases of the model. For example, convolutional layers learn translation-invariant functions, which are especially helpful in computer vision tasks [19].

In this paper we use a multi-layer perceptron (MLP) with a residual connection as the model architecture,

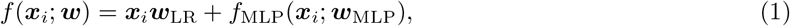

where the width and depth of *f*_MLP_(·) can be fixed beforehand or cross-validated. The residual network is essentially combining a linear regression with an MLP, which makes the neural network easier to train [22]. Moreover, it has been shown that residual networks are usually better than models without residual connections in genetics applications [8,29,30], and especially suitable for genotype-phenotype prediction, where linear main effects of the genotype often explain most of the phenotype variance.

### 3.2 Main effect prior (MEP)

We introduce the main effect prior as a novel regularization method for training neural networks. In the standard training procedure of neural networks, we find the optimal model parameters ***ŵ*** by minimizing a loss function, which consists of the average prediction error, 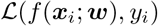, on the training set and a regularization function *Ω*(***w***) on the model parameters ***w*** to avoid overfitting,

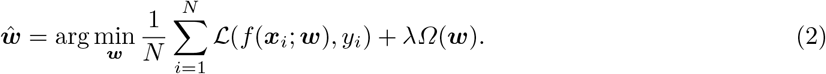

Here a cross-validated hyper-parameter *λ* on the training set controls the regularization strength. In the explanation prior framework [42,43,51], another regularization function, *Ω*′(***ϕ***(***w***), ***ϕ***\*), measuring the difference between the feature importance of the model and prior feature importance, is added:

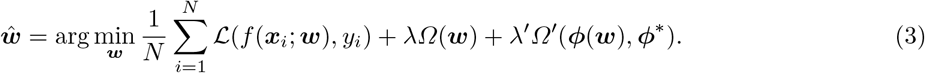

Here ***ϕ***(***w***) represents the feature importance of the neural network *f* (·; ***w***), and ***ϕ***\* is the prior feature importance provided by the user.

The published GWAS summary statistics are essentially the linear regression coefficients of SNPs when predicting the phenotype variable using each SNP independently. In order to derive priors from GWAS summary statistics, we calculate the multivariate linear regression which best approximates the neural network, and regularize its coefficients according to the explanation prior framework (Equation 3) towards the joint effects of SNPs, which can be derived from the published GWAS summary statistics (see Section 3.3). Specifically, we define the feature importance vector ***ϕ***(***w***) as the coefficient vector of a multiple linear regression that predicts *f* (*X*; ***w***), i.e., the projection of neural network predictions on the *X*:

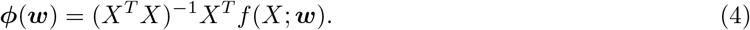

Assuming we have the prior joint main effects and corresponding standard deviations of SNPs, i.e., 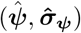, derived from the GWAS summary statistics of an external dataset, we define the main effect prior (MEP) as

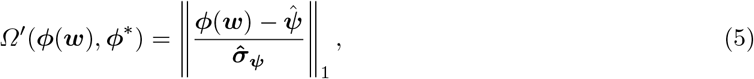

and the regularization coefficient *λ*′ is cross-validated in the same way as *λ*. Note that Equation 5 defines an *implicit* prior over neural network functions, instead of an *explicit* prior over the weights, in a fashion similar to the common weight decay regularization, which corresponds to the Gaussian prior. In our case, this prior encodes the inductive bias that the main effects implied by the neural network model should comply with the published summary statistics from the external data. In essence, our goal is to approximate the neural network with linear regression to provide importance weights matching earlier summary statistics, a priori. In this work, we focus on continuous phenotype prediction problems, so the prediction error 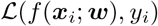 is measured with the squared error. We use *L*_1_ regularization to avoid overfitting, analogously to the LassoNet [29], i.e., *Ω*(***w***) = ||***w***||_1_, because the genotype matrix is potentially high-dimensional.

### 3.3 Constructing the prior

In this section, we provide details of constructing the prior joint main effects, i.e., 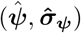 in Equation 5, from existing GWAS summary statistics.

#### Estimating the prior joint effects from marginal effects

In GWAS, the reported summary statistics are usually marginal effects of SNPs that are tested one by one with single-SNP models, which do not take the correlation between SNPs (i.e., the linkage disequilibrium) into account. In order to use the MEP defined in Equation 5, we first convert the marginal effects from GWAS summary statistics into joint effects.

We apply the earlier-studied approach [54] to estimate the joint effects of multiple SNPs from corresponding marginal effects without accessing the individual-level data. If the individual-level genotype-phenotype data is available, we could estimate the joint effects 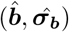 and marginal effects 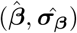 of multiple SNPs with the least-squares approach

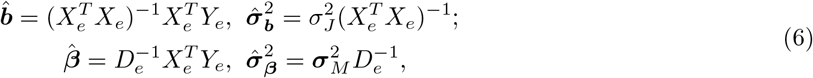

where (*X_e_, Y_e_*) represents the external genotype-phenotype data with size *N_e_* where the summary statistics have been estimated, *D_e_* is the diagonal matrix with entries from the diagonal of 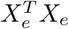, and 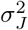 and 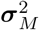 are the residual variances of the multi-SNP linear model and multiple single-SNP linear models. We assume that the mean of each SNP has been removed for simplicity.

From Equation 6, we can write the relations between 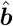 and 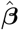 immediately [54]:

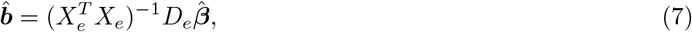

where 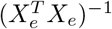 and *D_e_* are unavailable to us because individual-level data have not been published. However, assuming Hardy-Weinberg equilibrium (HWE) and knowing the allele frequency *f_j_* of SNP *j*, the *j*th element of *D_e_*_(*j*)_ be estimated by *D_e_*_(*j*)_ = 2*f_j_*(1 − *f_j_*)*N_e_*. Moreover, knowing the LD matrix *Σ_e_* of the external dataset, we can estimate 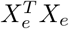 with

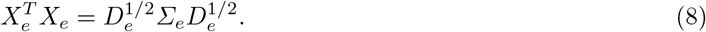

When *Σ_e_* is unavailable, the LD matrix from a reference sample (e.g., the training data) can be used instead.

Following the definition of residual variances of single and multiple linear regressions, we have the following relations between 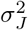 and 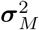:

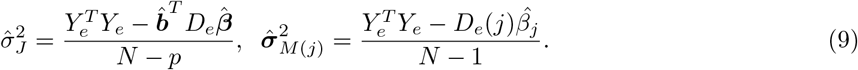

Hence, we can compute 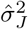 by first estimating 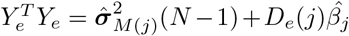. Then the standard deviation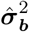 can be estimated with Equation 6 [54]. There are sophisticated approaches based on these ideas, such as LDpred [49] and PRS-CS [15], which additionally accommodate different effect size distributions within the Bayesian framework, and which can be applied to reconstruct the prior joint effects from GWAS summary statistics. In this work, we adopt the straightforward idea presented above and leave the exploration of the possible further enhancements for future work.

#### Calibrating the prior joint effects on the training set

After obtaining the prior joint effects 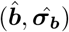 from external GWAS summary statistics, we propose to calibrate the prior joint effects on a target cohort. This is needed for example because the phenotype might have been scaled differently in different cohorts due to different measurement standards [24,35,36]. Moreover, it is necessary to normalize the phenotype of the training data to improve the neural network training. For these reasons, the raw prior joint effects usually reflect the actual influences on the target phenotype only up to a scaling factor.

We use a simple linear regression on the training set to calibrate the prior joint effect, by scaling the phenotype predicted with summary statistics 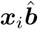 according to the target phenotype,

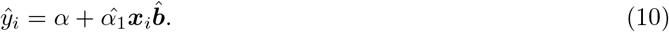

Thus, the calibrated prior joint effects that will be used in Equation 5 are given by 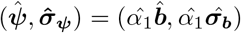. In Section 4.2, we show that this calibration is necessary especially when the external summary statistics are available from a different cohort than the target cohort.

## 4 Experiments

In this section, we first analyze multiple simulated data sets with different configurations to demonstrate the benefits of MEP. Then we apply the approach to two real-world metabolite data sets. Lastly, we conduct several ablation studies and a sensitivity analysis on the proposed method.

### 4.1 Simulation study

#### Setting

To construct the genotype dataset *X* for the simulation study, we select all SNPs (after basic quality control) from different numbers *p_g_* of genes associated with several cholesterol phenotypes from the UK Biobank [5], resulting in *p* SNPs in total. We simulate the phenotype *Y* by first simulating the “gene expression level” *G_g_* of each gene *g* according to a linear function of all SNPs *X*_(*g*)_ in that gene, and then generate phenotypes *Y* with main and interaction effects of *G_g_*:

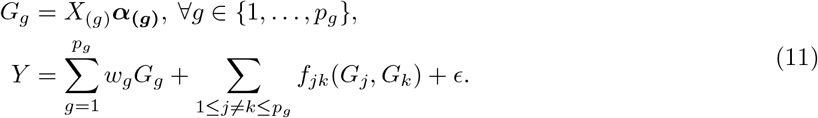

In the simulator, *f_jk_*(·, ·) represents the interaction function between genes *j* and *k*, chosen from three types: multiplicative, *G_j_* × *G_k_*, maximum, max{*G_j_, G_k_*}, and squared difference, (*G_j_* − *G_k_*)^2^. We set the parameters related to the main effect, i.e., ***α*_(*g*)_**and *w_g_*, such that the main effects explain ~90% of the signal variance, and we tune the variance of noise *E* to vary the signal-to-noise ratio (S/N) of the data. For each simulation, we randomly select 100,000 individuals to construct the external summary statistics. From the remaining individuals, we select *N* in the training set and 20,000 in the test set; hence, this corresponds to the simple setup where the summary statistics are from the same cohort as the target data. We study the performance of the MEP for different numbers of training data *N*, numbers of SNPs *p*, and signal-to-noise ratios (S/N). The two-stage generative process in Equation 11 reflects the idealized domain knowledge that SNPs within a gene affect how the gene is expressed, and the combined expression of multiple genes affects the phenotype. Furthermore, it reduces the complexity of simulating pairwise interactions, from *p*(*p* − 1)/2 to *p_g_*(*p_g_* − 1)/2 interactions. However, unlike [10], we do not assume this hierarchical structure of the simulator in the neural network that is used for prediction.

#### Results

We show the results of the simulation study with different settings in Figure 2. In general, we observe that neural networks with MEP, `NN+MEP (external)`, that incorporate the summary statistics from the external set outperform neural networks without MEP `(NN)` and linear regression `(LR)`, consistently in all settings. Moreover, we see in the middle panel that as more individual-level training data are available, the performance of `NN+MEP(external)` and NN becomes indistinguishable (e.g., when N=80k). In the last panel of Figure 2, we study how the number of SNPs affects the results. We notice that although the performance of all methods decreases as the number of SNPs increases (while keeping the total S/N fixed), the gap between `NN+MEP(external)` and NN increases. This is because models without MEP fail to learn the effects of high-dimensional SNPs from insufficient training data. Since the main effects of SNPs have already been captured by GWAS summary statistics on the large external dataset in `MEP(external)`, using MEP is especially beneficial for high-dimensional data.

**Fig. 2.**
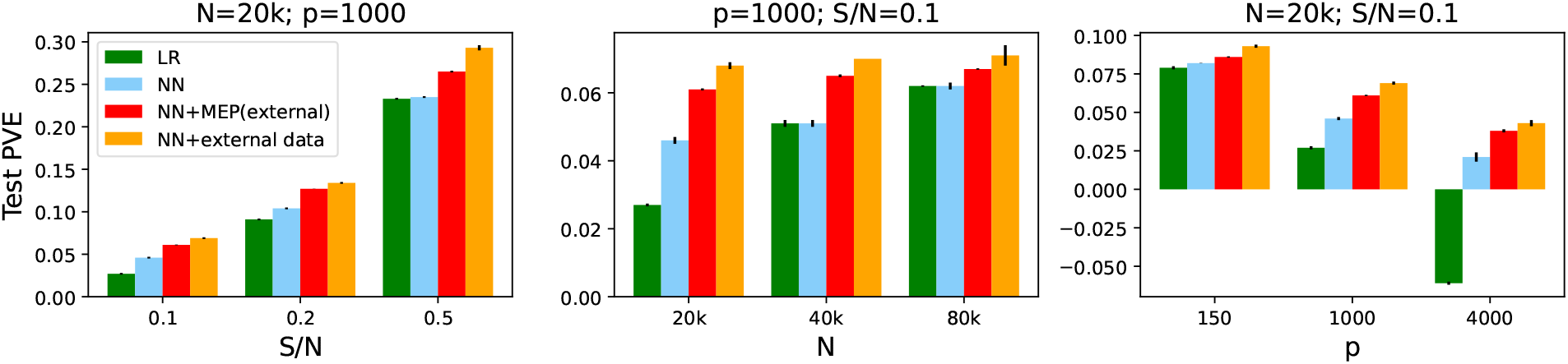
Results of the simulation study. Each bar shows the averaged test PVE (over 50 repeated experiments), and the error bar is the corresponding 95% CI. An unrealistic upper bound `(NN+external data)` is given by assuming that the individual-level external dataset is available, instead of only the GWAS summary statistics. With MEP that incorporates the GWAS summary statistics from the external dataset, `NN+MEP(external)` outperforms models trained on the target dataset without MEP in all settings.

The figure also plots results of the “golden standard” baseline constructed by training NNs on both the training set and the external set (`NN+external data`). This assumes that we can access the individual-level data of the large external cohort instead of only the GWAS summary statistics, and is hence a non-realistic upper bound of results. As expected, `NN+external data` results are better than all the realistic methods including `NN+MEP(external)` because the external dataset contains more information than its summary statistics. However, the performance gaps are relatively small when S/N is small (e.g., S/N=0.1 in the left panel), where the linear effects captured by the GWAS summary statistics are likely to explain most of the signal.

### 4.2 Real-world datasets

In this section, we consider the prediction of six different NMR metabolite phenotypes determined at Nightingale Health [53]: cholesterol and triglycerides in HDL, LDL, and VLDL, available in two real-world datasets UK Biobank [5] and FINRISK [4]. We use these two data sets to construct two kinds of realistic test cases: 1) the *homogeneous* case, where the summary statistics and the target dataset are both from different subsets of the same cohort, and 2) the *heterogeneous* case where the summary statistics are from an external cohort that is different from the cohort of the target data. In both cases, we use a large subset of the UK Biobank as the external data from which the summary statistics are extracted, and either a small subset of the UK Biobank or the FINRISK as the target data set in the homogeneous and heterogeneous cases, respectively.

#### Setting

To construct the homogeneous case study, we randomly split the UK Biobank dataset, which consists of 117, 981 individuals with fully measured NMR metabolomics, into two parts: an external dataset with 110, 000 individuals for which we estimate the GWAS summary statistics with PLINK1.9 [6], and a target dataset with 7,981 individuals, to train and test the prediction models. The heterogeneous case is exactly the same, except that we use the whole FINRISK (DILGOM07 more specifically) dataset, with 4,620 individuals, as the target data set (See Figure 1).

As suggested in [2,18], we reduce the dimensionality of the genotype in the target dataset and thus reduce the training complexity of the neural network, by applying LD clumping, which removes strongly correlated SNPs based on the GWAS summary statistics in the external dataset. The numbers of remaining SNPs for each phenotype are shown in Table 2 to 5. Moreover, we further test that our conclusions are not sensitive to the p-value threshold in LD clumping that determines the number of selected SNPs (See Section 4.4). For each experiment, we randomly split the target data into training (80%) and test (20%) sets. The genotypes are mean-centered and phenotypes standardized as part of data preprocessing. The hyper-parameters of each method are chosen via 5-fold cross-validation on the training set. We repeat each experiment with 10 different splits to evaluate the significance, and we measure the performance with the proportion of variance explained (PVE), i.e., *R*^2^, on the test set.

#### Baselines

We consider 9 different approaches: 1. linear regression `(LR)`, specifically BLUP [3]; 2. ridge regression `(Ridge)`, a `LR` with *L*_2_ regularization; 3. lasso regression `(Lasso)`, a `LR` with *L*_1_ regularization [28]; 4. elastic net, a `LR` with a combination of *L*_1_ and *L*_2_ regularization [20]; 5. gradient boosting tree `(XGBoost)`, implemented with XGBoost [9]; 6. neural networks `(NN)`, a 2-hidden-layer MLP with a residual connection discussed in Section 3.1; 7. the joint effect model derived from external GWAS (Section 3.3); 8. linear regression with MEP (`LR+MEP(external)`), i.e., replacing the neural network *f* (***x_i_***; *y_i_*) in Equation 3 with a linear regression; 9. neural networks with MEP (`NN+MEP(external)`).

#### Results

We first study whether the calibration of the main effects proposed in Equation 10 is necessary. In Table 1, we show the predictive performance, in terms of the test PVE, of the joint MEP before and after the calibration on the training set. We see that even in the simple <u>homogeneous</u> case, where the summary statistics and the target data are both from the UK Biobank, the calibration step improves the prediction in 3 out of 6 phenotype variables. Moreover, in the <u>heterogeneous</u> case, where the target data set is instead from the FINRISK, the calibration significantly improves the prediction for all phenotypes considered.

**Table 1.** Results of the calibration experiment. Averaged test PVE (over 10 random splits) of the joint effect prior before and after the calibration step on each phenotype in the homogeneous (UK Biobank, UKB) and heterogeneous (FINRISK,FIN) cases. The best result in each experiment is boldfaced, and the corresponding statistical significance is marked as stars. We observe that the calibration step is especially beneficial in the FINRISK dataset (heterogeneous cases).

| Priors | HDL.C |  | LDL.C |  | VLDL.C |  |
| --- | --- | --- | --- | --- | --- | --- |
|  | UKB | FIN | UKB | FIN | UKB | FIN |
| Joint effects | **0.078** | 0.052 | 0.060 | 0.048 | 0.064 | 0.004 |
| Calibrated joint effects | **0.078** | **0.055\*\*\*** | **0.064\*\*\*** | **0.050\*** | **0.065** | **0.026\*\*\*** |
|  | HDL.TG |  | LDL.TG |  | VLDL.TG |  |
|  | UKB | FIN | UKB | FIN | UKB | FIN |
| Joint effects | 0.074 | 0.023 | 0.084 | 0.032 | **0.063** | 0.034 |
| Calibrated joint effects | **0.075\*** | **0.039\*\*\*** | **0.086\*** | **0.040\*\*\*** | **0.063** | **0.036\*\*\*** |

<sup>1</sup> \*, \*\*, \*\*\* represent the p-value of corresponding test < 0.05, 0.01, 0.001 respectively.

**Table 2.**
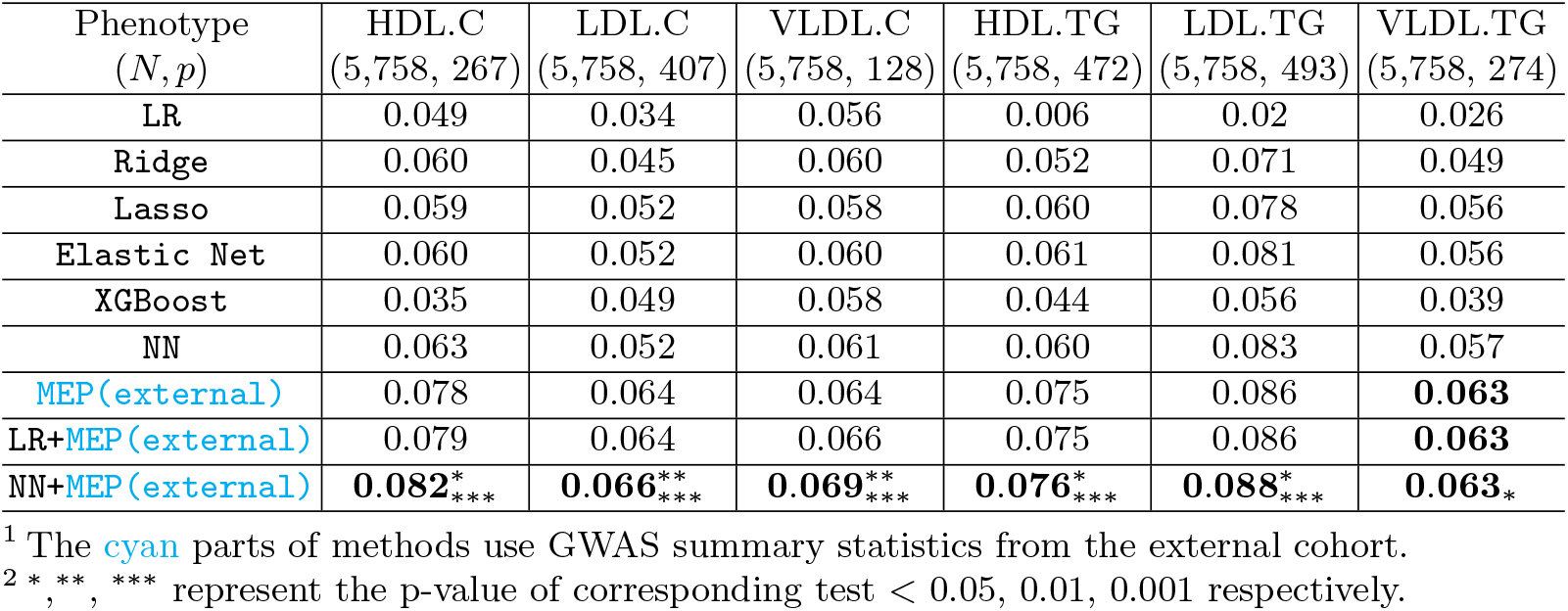
Comparison of the methods in the homogeneous case. Averaged test PVE (over 10 random splits) of all baseline methods on the target dataset (a subset of the UK Biobank). The dimension (*p*) and size of the training set (*N*) are shown for each phenotype. The best result in each experiment is boldfaced. We show the statistical significance of whether `NN+MEP(external)` is better than the second-best, with stars on the superscript, and better than without the MEP `(NN)`, with stars on the subscript.

Next, we compare the predictive performance of the different methods in the homogeneous and heterogeneous cases in Tables 2 and 3, respectively. First, we see that our proposed `NN+MEP(external)` has the (shared) highest accuracy for all 6 phenotypes in both cases. To check whether the improvement is statistically significant, we conduct two one-sided t-tests on the differences between test PVEs over 10 random splits to confirm: 1. whether the best model (`NN+MEP(external)`) is significantly better than the second-best model, and 2. whether `NN+MEP(external)` is significantly better than `NN`, i.e., the same model but without the MEP from the external dataset. The results of these two tests are shown in Tables 2 and 3 using stars in the superscript or subscript, respectively.

**Table 3.**
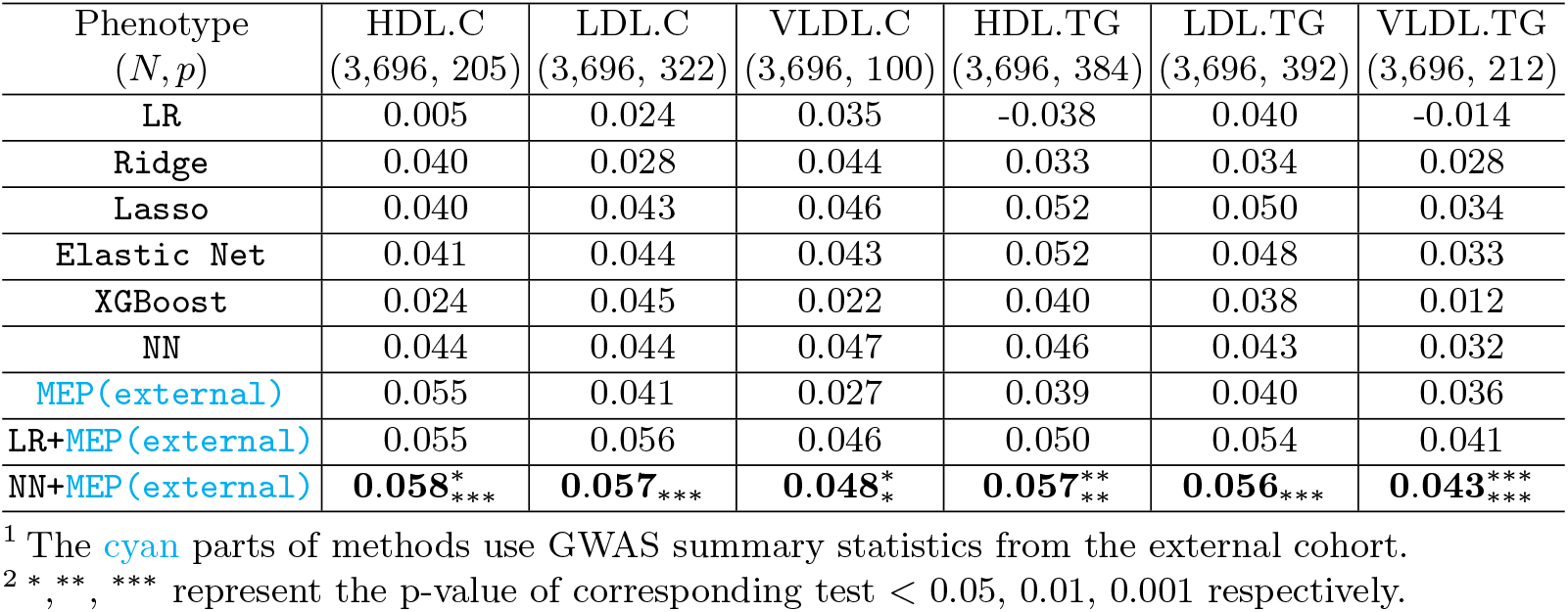
Comparison of the methods in the heterogeneous case. Averaged test PVE (over 10 random splits) of all baseline methods on the target dataset (FINRISK). The dimension (*p*) and size of the training set (*N*) are shown for each phenotype. The best result in each experiment is boldfaced. We show the statistical significance of whether `NN+MEP(external)` is better than the second-best, with stars on the superscript, and better than without the MEP `(NN)`, with stars on the subscript.

Specifically to the homogeneous case (Table 2), we see that using the `MEP (external)` as such (i.e., just using the model trained with the external summary statistics but without any adjustment with the target data) is already better than all baselines trained on the training data without the MEP. This is because the size of the external dataset is much larger than the training data and the effect sizes are homogeneous. However, even in this homogeneous case, information from the training set is helpful, as `LR+MEP(external)` and `NN+MEP(external)` improve the prediction performance further. Furthermore, `NN+MEP(external)` is significantly better than `LR+MEP(external)` in 5 out of 6 phenotypes, which shows the effect of incorporating nonlinear main effects and interactions in the prediction model.

In the more challenging heterogeneous case (Table 3), we notice that, different from the homogeneous case, `MEP(external)` as such does not always perform better than models trained on the small training set, e.g., on phenotypes VLDL.C, HDL.TG, and LDL.TG, even when the main effects are calibrated. This is an instance of negative transfer: if the external cohort is very different, findings from it may not generalize to the target cohort. This could be handled with hierarchical modeling, but doing so accurately would require access to the individual-level data instead of mere publicly available summary statistics. Consequently, when both training data from the target cohort and summary statistics from the external cohort are available, `NN+MEP(external)` achieves the significantly best prediction performance in 4 out of 6 phenotypes, and is always significantly better than not including the external MEP.

### 4.3 Ablation studies

In this section, we analyze the MEP in detail with several ablation studies.

We first study if using the joint effects of summary statistics (in Section 3.3) is significantly better than using the marginal effects, i.e., effects estimated separately for each SNP, as the prior to regularize the model `(NN+MEP(marginal)`). Instead of defining the feature importance in Equation 4 using the joint effects, we use the marginal effects of neural network predictions, such that

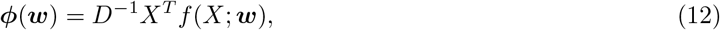

where *D* is the diagonal matrix with entries from the diagonal of *X^T^ X*. Moreover, we use the raw marginal effects 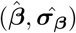 from GWAS to construct the prior instead of the derived joint prior in the MEP:

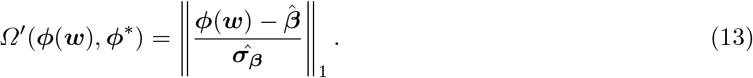

We show the performance of `NN+MEP(marginal)` in Table 4. We see that although using marginal effects as the prior improves prediction, for instance, in the phenotypes LDL.TG and VLDL.TG, the improvement is rather moderate compared with using the joint effects. This is because the correlation structure of SNPs is ignored in marginal effects, and they are especially important in phenotype prediction tasks [54].

**Table 4.** Results of the ablation studies. The studies are based on the heterogeneous setup (i.e. FINRISK as the target datasets). The performance is measured by the average test PVE over 10 random splits. The dimension (*p*) and size of the training set (*N*) are shown for each phenotype. We observe that using the marginal effects as prior `(NN+MEP(marginal)`) only has moderate improvement compared with `NN+MEP(external)`. Moreover, removing the standard deviation term from MEP (i.e., `NN+MEP-STD`) does not affect the results significantly.

| Phenotype<br>($N, p$) | HDL.C<br>(3,696, 205) | LDL.C<br>(3,696, 322) | VLDL.C<br>(3,696, 100) | HDL.TG<br>(3,696, 384) | LDL.TG<br>(3,696, 392) | VLDL.TG<br>(3,696, 212) |
| --- | --- | --- | --- | --- | --- | --- |
| NN | 0.044 | 0.044 | 0.047 | 0.046 | 0.043 | 0.032 |
| NN+MEP(marginal) | 0.044 | 0.045 | **0.048** | 0.050 | 0.055 | 0.037 |
| NN+MEP-STD | 0.059 | 0.051 | 0.046 | 0.051 | 0.059 | 0.042 |
| NN+MEP(empirical) | 0.045 | 0.044 | 0.046 | 0.046 | 0.044 | 0.032 |
| NN+MEP(external) | **0.061** | **0.056** | **0.048** | **0.057** | **0.060** | **0.043** |

<sup>1</sup> The cyan parts of methods use GWAS summary statistics from the external cohort.

We then study the effect of removing the standard deviation 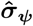 denominator from the MEP, i.e., using

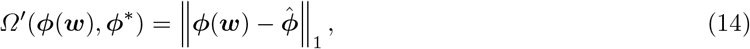

because the standard deviation term is often not provided in the more sophisticated constructions of the joint effects, such as LDpred [49] and PRS-CS [15]. From Table 4, we observe that `NN+MEP-STD` is still much better than `NN` without any MEP. Although `NN+MEP-STD` is worse than `NN+MEP(external)`, the difference is rather small in some phenotypes such as LDL.TG and VLDL.TG. Therefore, MEP is still applicable even the standard deviation term is not available.

Lastly, we study a heuristic empirical approach, where the joint effects are estimated using linear regression in the training dataset in the target cohort `NN+MEP(empirical)`, instead of on the external dataset. This corresponds to the situation where not even the GWAS summary statistics from another dataset are available. As expected, the test PVE of `NN+MEP(empirical)` is close to that of the `NN`, because no additional information is incorporated.

### 4.4 Sensitivity analysis

In Table 5, we analyze how sensitive the performance of each method is w.r.t. the p-value threshold in the LD clumping step, which affects the number of selected SNPs, on the HDL.C phenotype in FINRISK (the heterogeneous setting).

**Table 5.** **Results of the sensitivity study** w.r.t. the p-value threshold (per column) in LD clumping. The study is based on the heterogeneous setup (the HDL.C phenotype in FINRISK). The performance is measured by the average test PVE over 10 random splits. The dimension (*p*) and size of the training set (*N*) are shown for each p-value threshold. We show the statistical significance of whether `NN+MEP(external)` is better than the second-best, with stars on the superscript, and better than without the MEP `(NN)`, with stars on the subscript.

| p-value threshold<br>($N, p$) | 5e-6<br>(3,696, 676) | 5e-7<br>(3,696, 342) | 5e-8<br>(3,696, 205) | 5e-9<br>(3,696, 130) | 5e-10<br>(3,696, 108) |
| --- | --- | --- | --- | --- | --- |
| LR | -0.040 | -0.026 | 0.005 | 0.024 | 0.030 |
| Ridge | 0.035 | 0.036 | 0.040 | 0.043 | 0.043 |
| Lasso | 0.036 | 0.037 | 0.040 | 0.043 | 0.044 |
| Elastic Net | 0.036 | 0.037 | 0.041 | 0.043 | 0.043 |
| XGBoost | 0.023 | 0.023 | 0.024 | 0.024 | 0.024 |
| NN | 0.037 | 0.042 | 0.044 | 0.044 | 0.044 |
| MEP(external) | 0.063 | 0.061 | 0.055 | 0.051 | 0.048 |
| LR+MEP(external) | 0.063 | 0.060 | 0.055 | 0.052 | **0.052** |
| NN+MEP(external) | **0.065**<sup>*</sup><sub>***</sub> | **0.062**<sub>***</sub> | **0.058**<sup>*</sup><sub>***</sub> | **0.054**<sup>*</sup><sub>***</sub> | **0.052**<sub>***</sub> |

<sup>1</sup> The cyan parts of methods use GWAS summary statistics from the external cohort.
<sup>2</sup> \*, \*\*, \*\*\* represent the p-value of corresponding test < 0.05, 0.01, 0.001 respectively.

We observe that as we increase the p-value threshold, i.e., select more SNPs, the performance of the models without the MEP decreases (even achieve negative PVEs due to the overfitting), because they fail to learn the information in the additional SNPs from insufficient training data. However, the models trained with MEP achieve better performance with more SNPs, because the information from additional SNPs has already been captured by the GWAS summary statistics of the large cohort with `MEP(external)`. Moreover, the neural network with the MEP, `NN+MEP(external)`, consistently outperforms others.

## 5 Conclusion and discussion

In this work, we focused on genotype-phenotype prediction where a small individual-level target dataset and the GWAS summary statistics from a large external dataset were available. Due to the limited amount of data, even linear regression trained on the target dataset alone often failed to generalize well, not to mention complex models such as neural networks. We introduced a main effect prior for leveraging the GWAS summary statistics from an external dataset, which are commonly available, unlike additional individual-level data. Calculating the main effect prior only requires model predictions and input features; Thus, it is widely applicable with any machine learning model, although we focused on neural networks in this work. In both simulation studies and real-world datasets, we observed that neural networks could achieve significantly better prediction accuracy with the proposed main effect prior, compared with standard baselines, even on small training datasets.

Neural networks have many attractive properties in phenotype prediction, such as their ability to model arbitrary nonlinear main effects and feature interactions, but some challenges remain for future work. First, neural networks require massive individual-level training data, which are often unavailable due to privacy considerations. Here, we proposed a tractable solution by accessing the summary statistics from another large study. However, this may not be optimal because nonlinear effects and interactions in the external set, which could potentially benefit neural networks a lot, are lost when forming standard GWAS summary statistics. Recent advances in simulating and sharing individual-level data with differential privacy [23,46] could be a possible solution. Second, it would be interesting to devise model architectures that jointly incorporate the whole genome, instead of a subset of SNPs, to predict the phenotype. Unlike linear regression or summary statistics approaches, the number of parameters required to learn in a neural network is much larger than its number of input features. Moreover, we found that the relatively simple neural network architecture used here, i.e., a fully connected MLP with residual connections, was computationally heavy for wholegenome prediction. We resolved this with the two-stage approach whereby the neural network was built on genes selected previously by other methods. New architectures, including convolutional layers [2], that automatically achieve the required sparsity and parameter sharing would thus be needed for whole-genome studies. Moreover, Bayesian approaches, e.g., PRS-CS [15], designed specifically for estimating the joint genome-wide SNP effects from their summary statistics, would be particularly beneficial for constructing the MEP. Lastly, the proposed main effect prior could be generalized to classification settings, which cover some of the attractive phenotypes, e.g., different diseases, and we leave this as another interesting future direction.

## Acknowledgement

This work was supported by the Academy of Finland (Flagship programme: Finnish Center for Artificial Intelligence, FCAI, grants 319264, 292334, 286607, 294015, 336033, 315896, 341763), and EU Horizon 2020 (INTERVENE, grant no. 101016775). We also acknowledge the computational resources provided by the Aalto Science-IT Project from Computer Science IT.

